# A finite set of content-free pointers in visual working memory: MEG evidence

**DOI:** 10.1101/2024.07.26.605172

**Authors:** Xinchi Yu, Ellen Lau

## Abstract

Human visual working memory (VWM) is known to be capacity-limited, but the nature of this limit continues to be debated. Recent work has proposed that VWM is supported by a finite (∼ 3) set of content-free pointers, acting as stand-ins for individual objects and binding features together. Here, based on two visual working memory experiments (N=20 each) examining memory for simple and complex objects, we report a sustained MEG response over right posterior cortex whose magnitude tracks the core hypothesized properties of this system: load-dependent, capacity-limited and content-free. These results provide novel evidence for a finite set of content-free pointers underlying VWM.

## 1. Introduction

Visual working memory (VWM, or visual short-term memory, VSTM) for objects is a crucial yet extremely capacity-limited faculty for humans (Bays et al., 2024; Cowan, 2001; Fukuda et al., 2010; Luck & Vogel, 1997; Ma et al., 2014; Xu, 2017; Zhang & Luck, 2008). Recent work has attempted to reconcile earlier debates between “object-only” or “feature-only” theories of the capacity limit with theories of VWM architecture that contain elements of both and are thus better able to account for the ever-growing body of experimental findings at the same time (for a review see Ngiam, 2023). One such proposal is that VWM is implemented by a capacity-limited, content-free set of pointers (Swan & Wyble, 2014; Thyer et al., 2022; Xu & Chun, 2006; Yu & Lau, 2023): the finite set of pointers (or indexes/indexicals/tokens) act as stand-ins for each individual object in the mind (Balaban, Drew, et al., 2023; Balaban, Smith, et al., 2023; Pylyshyn, 1989), and bind features together for the corresponding objects. This model has allowed for separate capacity limits for pointers and visual features (Bowman & Wyble, 2007; Green & Quilty-Dunn, 2021; Hedayati et al., 2022; Swan & Wyble, 2014; Wang et al., 2017). Per theory, the representation for pointers in the brain, if any, needs to meet the following three criteria: (1) **load-dependent**: that this representation co-varies with (within-capacity) VWM load; (2) **capacity-limited (i.e**., **finiteness)**: that this representation reaches some kind of plateau or homeostasis beyond VWM capacity (of ∼ 3 objects); (3) **content-free**: the pointers contain only addresses, not the features themselves, and their function is to bind the addressed features together. The current study looks for such neural responses in MEG. Recent EEG (electroencephalography) evidence supports the existence of content-free representations in VWM with machine-learning multivariate decoding (Thyer et al., 2022) or univariate event-related potentials (Balaban et al., 2019; Balaban, Drew, et al., 2023; Wilson et al., 2012); however, it’s yet unclear whether these neural markers reflect the capacity limit. Furthermore, earlier fMRI (functional magnetic resonance imaging) evidence in fact suggested univariate representations in the brain that meets all three criteria, in regions including the posterior parietal cortex (Naughtin et al., 2016; Xu & Chun, 2006). The limitation of fMRI, however, is that it is hard to tell apart perceptual and VWM processes temporally; methods such as EEG and MEG (magnetoencephalography) have superior temporal resolution. Here we offer MEG evidence supporting the existence of VWM pointer representations that meet all three criteria: a right posterior MEG component is load-dependent, capacity-limited (finite) and content-free at the same time.

Previous MEG studies have identified load-dependent, capacity-limited responses during VWM in left and right posterior channels (Mitchell & Cusack, 2011; Robitaille et al., 2009, 2010), but with a bilateral-presentation paradigm^1^, with the hope of finding a CDA [contralateral delay activity]-like response as in EEG (which was not very successful). In the current study, we employed a similar logic as in the pioneering fMRI study of Xu & Chun (2006). In Experiment 1, we tested whether the left and right posterior effects in MEG are still load-dependent and capacity-limited with a central-presentation paradigm (as used in Xu & Chun, 2006). To preview, we found a reliable load-dependent and capacity-limited effect in right posterior channels. In Experiment 2, we showed that this right posterior load-dependent, capacity-limited effect is also content-free. Therefore, this right posterior activity (RPA) meets all three criteria for a neural index of VWM pointers.

## 2. Methods

### 2.1. Participants

For Experiment 1, 20 participants entered data analysis (12 female, age 19-41, *M*=24, 3 left-handed); 1 additional participant’s data was excluded from analysis due to excessive noise (see MEG analysis). For Experiment 2, 20 participants entered data analysis (13 female, age 18-37, *M*=21, 2 left-handed); 1 additional participant’s data was excluded from analysis due to excessive noise. All subjects reported having normal or corrected-to-normal vision. Informed consent was obtained from all participants and they received monetary reimbursement for their participation. Procedures were approved by the UMCP IRB Office.

### 2.2. Procedure

Experiments 1 and 2 had the same procedure. Each subject was presented with two blocks of 200 trials each, which differed in the features of the objects: one block in which the objects were colored squares, and another block in which the objects were complex shapes. The order of the blocks was balanced across subjects. For each block, there were 50 trials for each VWM load condition (1, 2, 4 and 6); within each 50 trials, half of the trials had a match probe and the other half had a change probe.

In each trial, the participants were first presented with a fixation cross with a duration of 500 ms (Experiment 1) or 800 ms (Experiment 2) against a grey background, followed by a sample screen with a duration of 160 ms (Experiment 1) or 200 ms (Experiment 2)^2^. Then, they were presented with the sample; see Figure 1. The sample consisted of 1, 2, 4, 6 different objects: for colored-square trials, they were sampled from 8 colors (black [0,0,0], dark blue [0,0,139], green [0,255,0], light blue [173,216,230], pink [255,192,203], red [255,255,0], white [255,255,255] and yellow [255,255,0]); for complex-shape trials, they were sampled from 9 complex shapes similar to those used in Xu & Chun (2006). 8 fixed locations were evenly distributed 4.5° from the center of the screen, and these locations were marked by 8 light grey squares (2.5° × 2.5° each). The objects (colored squares or complex shapes) were presented at the center of the light grey squares, spanning 2.1° × 2.1°. The light grey squares served as a visual cue to encourage the individuation of each object and to discourage grouping. A blank screen delay of 1000 ms followed the sample, after which the probe appeared on the screen. At the probe, only one object was presented at one of the 8 locations. On “match” trials, this object was the same one as the one in the sample for that location; on “change” trials, this probe object was randomly selected from other objects used in this block. Participants pressed one button to indicate “match”, and pressed another button to indicate “change” (buttons balanced across participants), with no time limit for response. After their response, they were shown a feedback for 600 ms, followed by an inter-trial interval (ITI) of 2500-3500 ms. During the ITI, a line of asterisks was presented on the screen; before the experiment, participants were instructed to blink only during this period, and to avoid blinking during the trials. While the fixation cross was presented on the screen, the participants were instructed to fixate at the center of the screen. They were also instructed against verbal rehearsal. At least 8 practice trials for each block were run before the main experiment for each participant, and a self-paced break was administered every 50 trials. Experiment 1 analyses included only the colored square block^3^, while Experiment 2 analyses included both a colored square block and a complex shape block (order balanced across participants). The experiment was run in MATLAB R2009a with Psychtoolbox 3 (Brainard, 1997; Kleiner et al., 2007).

**Figure 1.**
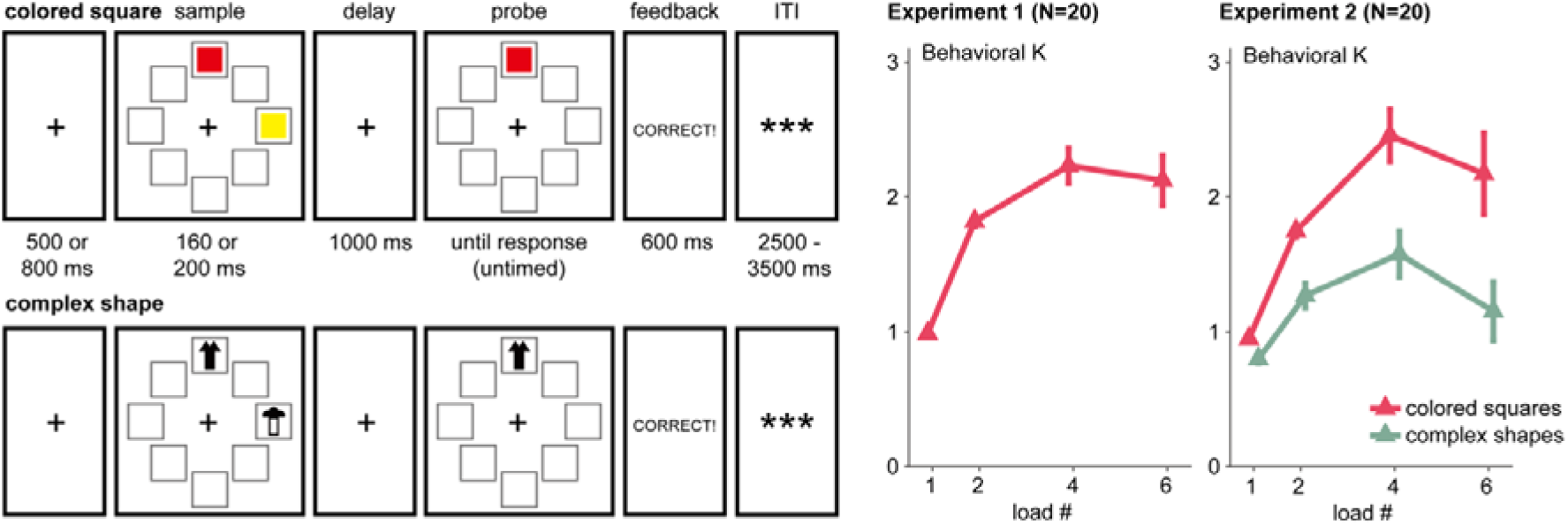
Experimental procedure and behavioral results (Cowan’s K) across Experiments 1 and 2. Error bars stand for standard error. ITI: inter-trial interval. For detailed presentation parameters of the stimuli see Procedure.

### 2.3. Behavioral analysis

Based on the hit rates and false alarm rates for each condition, we calculated Cowan’s K (Cowan, 2001; Rouder et al., 2011), which reflects the number of objects successfully held in VWM. The formula for Cowan’s K is as follows: K=N×(H-F), where N is sample load (i.e., 1, 2, 4 or 6), H is the hit rate for detecting the change in the probe on change trials, and F is the false alarm rate (erroneously “hallucinating” a change in the probe on match trials).

### 2.4. MEG recording

Prior to recording, five head position indicator coils were affixed to each participant’s head, and the position of these coils relative to the nasion and tragus, as well as the participant’s head-shape, was digitized using a Polhemus 3SPACE FASTRAK system in order to determine the participant’s accurate placement in the MEG dewar. During the experimental sessions, participants laid supine in a dimly lit magnetically shielded room (Yokogawa Industries, Tokyo, Japan). Continuous MEG recording was executed using a 160-channel axial gradiometer whole-head system (Kanazawa Institute of Technology, Kanazawa, Japan), and data was sampled at 1000 Hz (60 Hz online notch filter, DC-200 Hz recording bandwidth).

### 2.5. MEG analysis

MEG data was analyzed by customized code in MATLAB R2020b with MNE-Python 1.5.1 (Gramfort et al., 2013). First, noisy and flat channels were identified for each participant with Maxwell filtering, and these channels were interpolated with the MNE algorithm (no channels in the regions-of-interest described below required interpolation). Then the environmental magnetic interferences were suppressed using temporal signal space separation (tSSS; Taulu & Simola, 2006). A low-pass infinite impulse response (IIR) filter with an upper cutoff frequency of 40 Hz was applied to the MEG data. Ocular and cardiac artifacts were removed using independent component analysis (ICA; fastica algorithm); 2-5 ICA components were removed from each participant. Epochs of −200:1200ms time-locked to the presentation of sample stimuli in each trial were extracted. The 200 ms pre-stimulus interval was used as the baseline interval. Trials with a maximum peak-to-peak signal amplitude higher than 3000 fT were excluded. Participants with less than 40 valid trials per condition were excluded from data analysis (1 for each experiment, see Participants). For the analyzed participants, only 0.7% (Experiment 1), 0.1% (Experiment 2, colored squares), and 0.4% (Experiment 2, complex shapes) of all trials were excluded. For each participant and each condition (i.e., a certain load condition in a certain feature block), we calculated the mean event-related field (ERF) response for each channel.

The first 20 trials in each condition of Experiment 1 were used to determine regions-of-interest (ROIs) across both experiments, and as such were excluded from all subsequent analyses. The contrast used for ROI selection was the mean ERF difference between larger (load 4 and 6) vs. smaller (load 1 and 2) conditions (Figure 2). In line with previous reports (Mitchell & Cusack, 2011; Robitaille et al., 2009, 2010), we observed two opposing maxima in posterior channels. We took three channels each from the two maxima to construct right and left parietal ROIs (right: MEG054, MEG055, MEG071; left: MEG051, MEG050, MEG076), and computed mean ERFs for each from 400:1200 ms post-stimulus-onset to capture the VWM maintenance (as opposed to perception) period.

**Figure 2.**
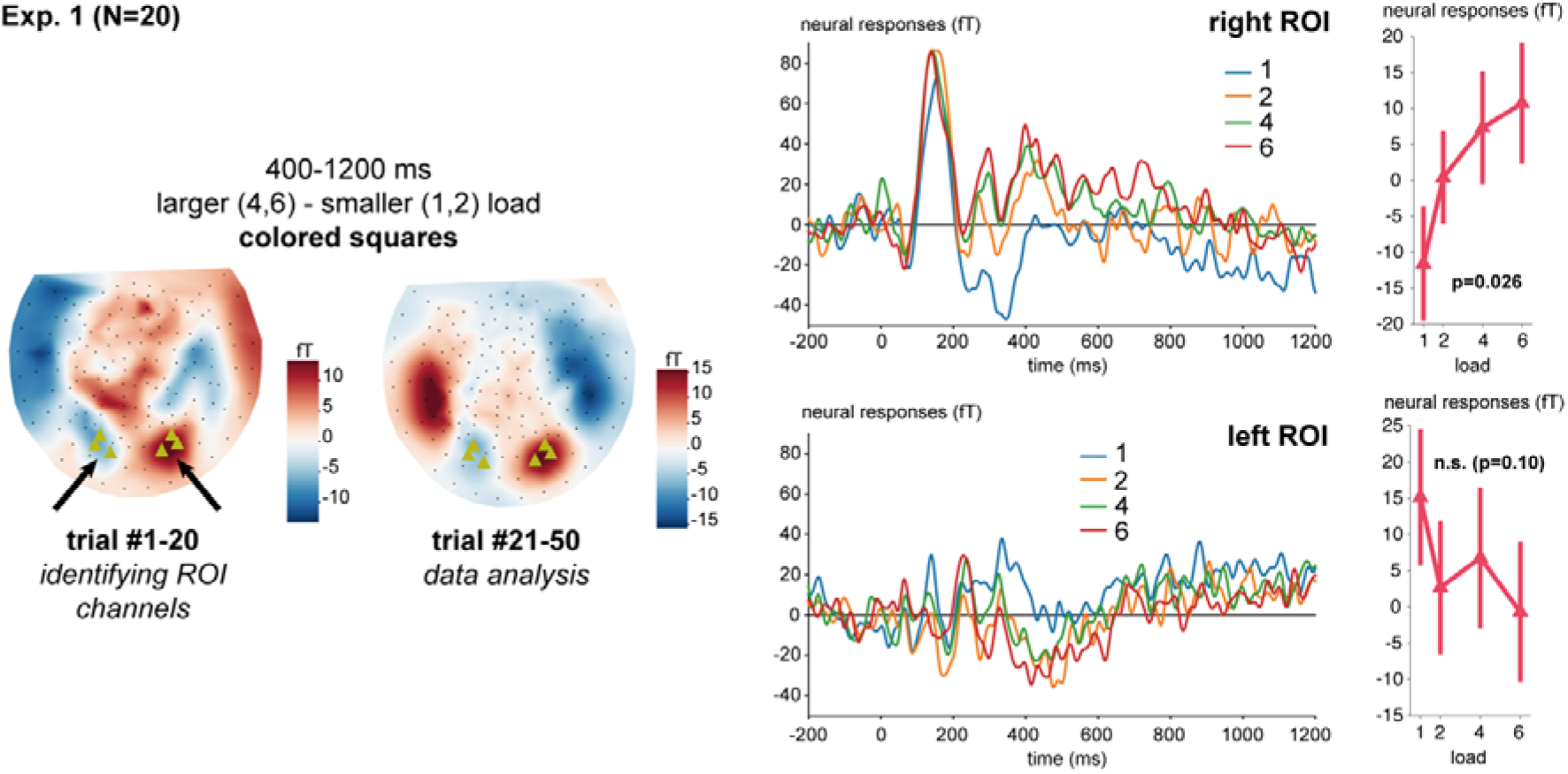
A summary of the MEG results in Experiment 1. Error bars stand for standard error. ROI: region of interest. The ROI channels are marked with green triangles.

Comparisons between conditions were statistically evaluated with repeated-measures ANOVA (using the Greenhouse-Geisser whenever the sphericity assumption was not met), and post hoc comparisons were Bonferroni-corrected. In order to better evaluate whether there was a response plateau beyond the capacity limit, we also administered Bayesian repeated-measures ANOVA with a uniform prior, and the post hoc comparisons were administered with Bayesian paired samples t test (with a default Cauchy prior). All statistical analyses reported in this paper were conducted with JASP 0.18.1 (JASP Team, 2023) and Bayesian factors (BFs) were interpreted based on van Doorn et al. (2019): a BF higher than 10 or lower than 1/10 is considered “strong”, between 3-10 or 1/10-1/3 is considered “moderate”, and between 1/3-3 is considered “weak”.

## 3. Results

### 3.1. Experiment 1

#### 3.1.1. Behavioral results

Repeated measures ANOVA revealed a significant main effect of load, *F*(1.80,34.28) = 30.22, *p* < 0.001. Post-hoc comparisons (Bonferroni corrected) revealed a significant difference between loads 1 vs. 2 (*t*(19) = −5.75, *p* < 0.001), 1 vs. 4 (*t*(19) = −8.59, *p* < 0.001), 1 vs. 6 (*t*(19) = −7.84, *p* < 0.001) and 2 vs. 4 (*t*(19) = −2.84, *p* = 0.038). The difference between 2 vs. 6 (*t*(19) = −2.10, *p* = 0.24) did not reach statistical significance, nor did the difference between 4 vs. 6 (*t*(19) = 0.74, *p* = 1). Bayesian repeated measures ANOVA revealed strong evidence for the effect of load, BF_incl_ = 1.66×10^10^. There was strong evidence for a difference between 1 vs. 2 (BF_10_ = 1.09×10^12^), 1 vs. 4 (BF_10_ = 1.73×10^6^), 1 vs. 6 (BF_10_ = 4.13×10^3^), and 2 vs. 4 (BF_10_ = 16.11), and moderate evidence against a difference between 4 vs. 6 (BF_10_ = 0.28), suggesting a plateau of Cowan’s K beyond a load of ∼ 4. The evidence for a difference between 2 vs. 6 was weak (BF_10_ = 0.79).

#### 3.1.2. MEG results

The repeated measures ANOVA for MEG responses in the right ROI (Figure 2) showed a statistically significant main effect of load, *F*(1.98,37.52) = 4.06, *p* = 0.026. Post-hoc comparisons (Bonferroni corrected) revealed a significant difference between loads 1 vs. 4 (*t*(19) = −2.73, *p* = 0.050) and 1 vs. 6 (*t*(19) = −3.23, *p* = 0.012). The other comparisons did not reach statistical significance (*p’s* > 0.5). Bayesian repeated measures ANOVA revealed moderate evidence for the effect of load, BF_incl_ = 4.15. There was moderate evidence for the difference between 1 vs. 6 (BF_10_ = 3.71), as well as moderate evidence against a difference between 4 vs. 6 (BF_10_ = 0.27). Evidence for other differences was weak (1/3 < BF_10_ < 3). We note that our number of trials per load condition (∼ 30) here is rather low compared to previous MEG studies of this kind (Robitaille et al., 2009: 384 trials/condition; Robitaille et al., 2010: 200 trials/condition; Mitchell & Cusack, 2011: 120 trials/condition) but with a similar sample size. Below we pool all 40 participants across the two experiments for a higher-powered analysis.

In the left ROI, the main effect of load was not statistically significant, *F*(3,57) = 2.15, *p* = 0.10. Evidence for the effect of load was also weak, BF_incl_ = 0.62.

### 3.2. Experiment 2

#### 3.2.1. Behavioral results

A repeated measures ANOVA crossing load and feature (colored square vs. complex shape), showed a significant main effect of load on Cowan’s K, (*F*(1.51,28.76) = 16.38, *p* < 0.001), a main effect of feature (*F*(1,19) = 50.45, *p* < 0.001), and a significant interaction (*F*(2.13,40.48) = 7.35, *p* = 0.002). Bayesian repeated measures ANOVA also revealed strong evidence for the effect of load (BF_incl_ = 2.42×10^7^), feature (BF_incl_ = 4.61×10^5^) and the interaction effect (BF_incl_ = 715.77). The main effect of feature reflected overall higher performance for colored squares than complex shapes, in line with previous observations (e.g., Xu & Chun, 2006).

We followed up the interaction with one-way ANOVAs at each level of feature. In colored square trials, the main effect of load was statistically significant, *F*(1.55,29.42) = 18.16, *p* < 0.001. Post-hoc comparisons (Bonferroni corrected) revealed significant differences between loads 1 vs. 2 (*t*(19) = −3.67, *p* = 0.003), 1 vs. 4 (*t*(19) = −6.92, *p* < 0.001), 1 vs. 6 (*t*(19) = −5.62, *p* < 0.001) and 2 vs. 4 (*t*(19) = −3.25, *p* = 0.012). The differences between 2 vs. 6 (*t*(19) = −1.95, *p* = 0.34) and 4 vs. 6 (*t*(19) = 1.30, *p* = 1) were not statistically significant. Bayesian repeated measures ANOVA also suggested strong evidence for the effect of load, BF_incl_ = 1.82×10^6^. There was strong evidence for the difference between 1 vs. 2 (BF_10_ = 1.36×10^11^), 1 vs. 4 (BF_10_ = 1.08×10^5^), 1 vs. 6 (BF_10_ = 63.91) and 2 vs. 4 (BF_10_ = 87.07). Evidence for the difference between 2 vs. 6 (BF_10_ = 0.61) and 4 vs. 6 (BF_10_ = 0.44) was weak.

In complex shape trials, the main effect of load was also significant (*F*(1.99,37.74)=7.01, *p*=0.003). Post-hoc comparisons (Bonferroni corrected) revealed significant differences between loads 1 vs. 2 (*t*(19) = −2.74, *p* = 0.049) and 1 vs. 4 (*t*(19) = −4.53, *p* < 0.001). The other differences were not statistically significant (*p’s* >0.09). Bayesian repeated measures ANOVA also suggested strong evidence for the effect of load, BF_incl_ = 93.00, and for the difference between 1 vs. 2 (BF_10_ = 6.03×10^4^) and 1 vs. 4 (BF_10_ = 136.50). Evidence for the other differences was weak (1/3 < BF_10_ < 3).

In sum, the colored square and complex shape comparisons patterned similarly in showing main effects of load. The interaction between feature and load appeared to be driven only by the slight drop observed in performance for complex shapes at load 6.

#### 3.2.2. MEG results

The 2 × 4 (feature × load) repeated measures ANOVA for MEG responses in the right ROI (Figure 3) showed a significant main effect of load (*F*(1.50,28.47) = 6.31, *p* = 0.010).

**Figure 3.**
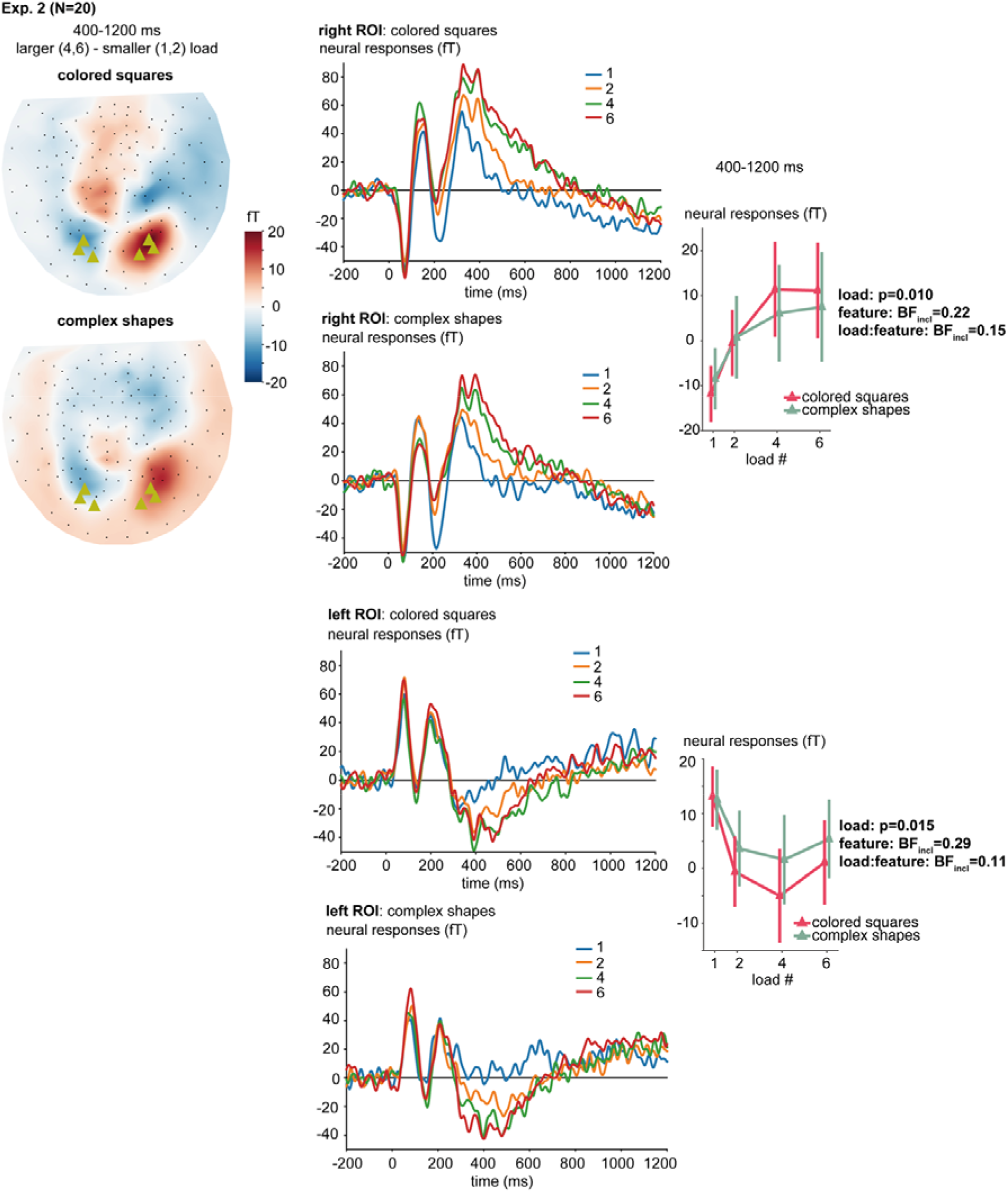
A summary of the MEG results in Experiment 2. Error bars stand for standard error. ROI: region of interest. The ROI channels are marked with green triangles.

The main effect of feature (*F*(1,19) = 0.086, *p* = 0.77) and the interaction effect (*F*(3,57) = 0.90, *p* = 0.45) were not significant. Post-hoc comparisons between load conditions (Bonferroni corrected) revealed a significant difference between loads 1 vs. 4 (*t*(19) = −3.68, *p* = 0.003) and 1 vs. 6 (*t*(19) = −3.79, *p* = 0.002); the other differences were not significant (*p’s* ≥ 0.30). Bayesian repeated measures ANOVA revealed moderate evidence against the inclusion of feature (BF_incl_ = 0.22) and the interaction effect (BF_incl_ = 0.15) in the model, and strong evidence for the inclusion of load (BF_incl_ = 29.78) in the model. There was moderate evidence for the difference between 1 vs. 2 (BF_10_ = 4.80) and 1 vs. 6 (BF_10_ = 29.58) and strong evidence for the difference between 1 vs. 4 (BF_10_ = 44.75); there was moderate evidence against the difference between 4 vs. 6 (BF_10_ = 0.17), suggesting a plateau beyond ∼ 4. Evidence for the difference between 2 vs. 4 (BF_10_ = 1.80) and 2 vs. 6 (BF_10_ = 2.18) was weak.

For the left ROI, the main effect of load was significant, (*F*(1.97,37.36) = 4.75, *p* = 0.015). The main effect of feature (*F*(1,19) = 1.37, *p* = 0.26) and the interaction effect (*F*(3,57) = 0.42, *p* = 0.74) were not statistically significant. Post-hoc comparisons (Bonferroni corrected) revealed a significant difference between loads 1 vs. 2 (*t*(19) = 2.79, *p* = 0.043) and 1 vs. 4 (*t*(19) = 3.58, *p* = 0.004); the other differences were not statistically significant (*p’s* > 0.12). Bayesian repeated measures ANOVA revealed moderate evidence against the inclusion of feature (BF_incl_ = 0.29) and the interaction effect (BF_incl_ = 0.11) in the model, and moderate evidence for the inclusion of load (BF_incl_ = 4.79) in the model. There was moderate evidence for the difference between 1 vs. 2 (BF_10_ = 22.98) and 1 vs. 4 (BF_10_ = 9.58), and moderate evidence against the difference between 2 vs. 4 (BF_10_ = 0.25) and 2 vs. 6 (BF_10_ = 0.19), suggesting a plateau beyond ∼ 2. Evidence for the difference between 1 vs. 6 (BF_10_ = 2.36) and 4 vs. 6 (BF_10_ = 0.79) was weak.

Pooling colored square data from all 40 participants across Experiments 1 and 2 together allowed us to conduct a high-powered analysis to determine whether the two ROIs exhibited the monotonic increase up to 4 followed by a plateau, as predicted if there is a limited pointer capacity of ∼3 (Figure 4). For the right ROI, the Bayesian paired-sample t-test revealed moderate evidence for differences between loads 1 vs. 2 (BF_10_ = 16.07), 1 vs. 4 (BF_10_ = 28.00), 2 vs. 6 (BF_10_ = 5.50) and strong evidence for a difference between 1 vs. 6 (BF_10_ = 134.59). There was also moderate evidence against a difference between 4 vs. 6 (BF_10_ = 0.19). Evidence for the difference between 2 vs. 4 was weak (BF_10_ = 2.18) but was still in the direction of a difference. Overall, this suggests a plateau beyond ∼ 4 items for the right ROI.

**Figure 4.**
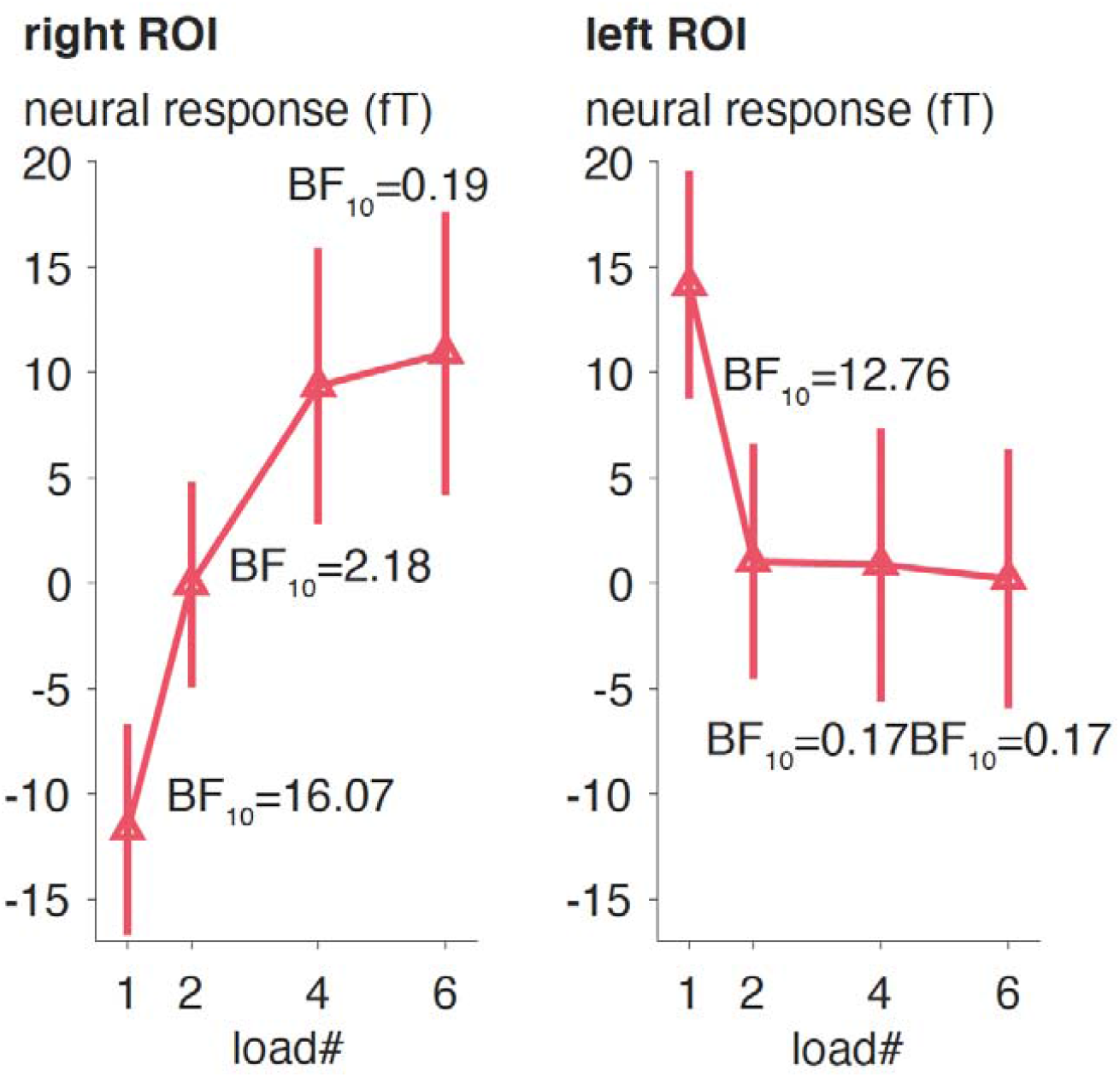
MEG responses during the VWM period (400:1200 ms) for the colored squares in the right and left ROIs, pooled across all 40 participants in Experiments 1 and 2. Error bars stand for standard error. ROI: region of interest.

For the left ROI, the Bayesian paired-sample t-test revealed strong evidence for a difference between loads 1 vs. 2 (BF_10_ = 12.76) and moderate evidence for a difference between 1 vs. 6 (BF_10_ = 7.94), and moderate evidence against a difference between loads 2 vs. 4 (BF_10_ = 0.17), 2 vs. 6 (BF_10_ = 0.17), and 4 vs. 6 (BF_10_ = 0.17). Evidence for the difference between 1 vs. 4 was weak (BF_10_ = 2.44) but was still in the direction of a difference. Overall, this suggests a plateau beyond ∼ 2 items for the left ROI.

## 4. Discussion

In this study, we observed univariate responses in MEG over right posterior sites (the right posterior activity, RPA) that met all three criteria for VWM pointers: load-dependent, capacity-limited, and content-free. This neural response changed monotonically as VWM load increased, reached a plateau at ∼ 4 objects (in line with the classic VWM capacity limit), and was insensitive to the difference in type and number of features bound to each object in the colored square vs. complex shape conditions even though these differences impacted behavioral performance. Given its maxima over right posterior channels, this response is likely to be of posterior parietal origin, in line with previous fMRI effects attributed to VWM pointers (Naughtin et al., 2016; Xu & Chun, 2006), and consistent with the involvement of the posterior parietal cortex in VWM maintenance in general (Marois & Todd, 2004; Matsuyoshi et al., 2010; Xu, 2023). This awaits to be confirmed by future studies with higher spatial-temporal resolution (e.g., combining MEG with structural MRI, intracranial recordings). Our current study serves as an important complement to prior fMRI studies, in that we confirmed VWM pointer-like responses during the VWM maintenance time window in particular. The right-lateralization of our current effects invites future neuroimaging research on the lateralization of VWM substrates. Intriguingly, this is in line with some neuropsychological reports suggesting that unilateral *right* posterior parietal lesions are sufficient for an impaired VWM (for a review and nuanced discussions see Olson & Berryhill, 2009).

We also found that left posterior channels exhibited a load-dependent, capacity-limited and content-free effect, but this response plateaued at only ∼ 2 objects. While this different response profile awaits future investigation, it could reflect cognitive effort related to pattern-separating more than one feature of the same type (e.g., when one needs to hold multiple colors in VWM) or mechanisms related to attention allocation (Todd & Marois, 2005).

## CRediT authorship contribution statement

**Xinchi Yu** Conceptualization, data curation, investigation, formal analysis, methodology, software, visualization, writing-original draft, writing-review & editing. **Ellen Lau:** Conceptualization, funding acquisition, methodology, supervision, writing-original draft, writing-review & editing.

## Data availability

Data is available upon request.

## Acknowledgements

We would like to thank Marco Chia-Ho Lai and Ciaran Stone for the discussions on MEG data analysis.

## Footnotes

1 In EEG studies examining the CDA response, separate stimuli are presented on the left and right sides of the screen respectively. The participants are cued beforehand to memorize the memorandum on one of the two sides. The CDA is computed by subtracting ipsilateral from contralateral ERP (event-related potential) responses during the delay, and its amplitude is load-dependent and capacity-limited (Luria et al., 2016; Vogel & Machizawa, 2004). However, most prior MEG studies were not successful in finding a CDA-like effect in MEG (likely due to the fact that MEG and EEG are not capturing exactly the same neural responses, Becke et al., 2015); rather, they observed a load-dependent, capacity-limited effect in the raw responses at left/right posterior channels.

2 Because of a programming issue, 5 participants in Experiment 1 had a fixation cross duration of 800 ms, and 5 participants in Experiment 2 had a sample duration of 160 ms.

3 We originally included a complex shape block in Experiment 1, but due to a timing error we were unable to analyze the data for those trials.

## References

Balaban, H., Drew, T., & Luria, R. (2019). Neural evidence for an object-based pointer system underlying working memory. Cortex, 119, 362–372. 10.1016/j.cortex.2019.05.008

Balaban, H., Drew, T., & Luria, R. (2023). Dissociable online integration processes in visual working memory. Cerebral Cortex, 33(23), 11420–11430. 10.1093/cercor/bhad378

Balaban, H., Smith, K., Tenenbaum, J. B., & Ullman, T. (2023). Electrophysiology reveals that intuitive physics guides visual tracking and working memory. PsyArXiv. 10.31234/osf.io/pr4ym

Bays, P. M., Schneegans, S., Ma, W. J., & Brady, T. F. (2024). Representation and computation in working memory. Nature Human Behaviour.

Becke, A., Müller, N., Vellage, A., Schoenfeld, M. A., & Hopf, J. M. (2015). Neural sources of visual working memory maintenance in human parietal and ventral extrastriate visual cortex. NeuroImage, 110, 78–86. 10.1016/j.neuroimage.2015.01.059

Bowman, H., & Wyble, B. (2007). The simultaneous type, serial token model of temporal attention and working memory. Psychological Review, 114(1), 38–70. 10.1037/0033-295X.114.1.38

Brainard, D. H. (1997). The Psychophysics Toolbox. Spatial Vision, 10(4), 433–436. 10.1163/156856897X00357

Cowan, N. (2001). The magical number 4 in short-term memory: A reconsideration of mental storage capacity. Behavioral and Brain Sciences, 24(1), 87–114. 10.1017/S0140525X01003922

Fukuda, K., Awh, E., & Vogel, E. K. (2010). Discrete capacity limits in visual working memory. Current Opinion in Neurobiology, 20(2), 177–182. 10.1016/j.conb.2010.03.005

Gramfort, A., Luessi, M., Larson, E., Engemann, D. A., Strohmeier, D., Brodbeck, C., Goj, R., Jas, M., Brooks, T., Parkkonen, L., & Hämäläinen, M. (2013). MEG and EEG data analysis with MNE-Python. Frontiers in Neuroscience, 7(7 DEC), 1–13. 10.3389/fnins.2013.00267

Green, E. J., & Quilty-Dunn, J. (2021). What Is an Object File? The British Journal for the Philosophy of Science, 72(3), 665–699. 10.1093/bjps/axx055

Hedayati, S., O’Donnell, R. E., & Wyble, B. (2022). A model of working memory for latent representations. Nature Human Behaviour, 6(5), 709–719. 10.1038/s41562-021-01264-9

JASP Team. (2023). JASP 0.18.1.

Kleiner, M., Brainard, D., Pelli, D., Ingling, A., Murray, R., & Broussard, C. (2007). ECVP ‘07 Abstracts. Perception, 36(1_suppl), 1–235. 10.1177/03010066070360S101

Luck, S. J., & Vogel, E. K. (1997). The capacity of visual working memory for features and conjunctions. Nature, 390(6657), 279–281. 10.1038/36846

Luria, R., Balaban, H., Awh, E., & Vogel, E. K. (2016). The contralateral delay activity as a neural measure of visual working memory. Neuroscience and Biobehavioral Reviews, 62, 100–108. 10.1016/j.neubiorev.2016.01.003

Ma, W. J., Husain, M., & Bays, P. M. (2014). Changing concepts of working memory. Nature Neuroscience, 17(3), 347–356. 10.1038/nn.3655

Marois, R., & Todd, J. J. (2004). Capacity limit of visual short-term memory in human posterior parietal cortex. Nature, 428, 751―754.

Matsuyoshi, D., Ikeda, T., Sawamoto, N., Kakigi, R., Fukuyama, H., & Osaka, N. (2010). Task-irrelevant memory load induces inattentional blindness without temporo-parietal suppression. Neuropsychologia, 48(10), 3094–3101. 10.1016/j.neuropsychologia.2010.06.021

Mitchell, D. J., & Cusack, R. (2011). The temporal evolution of electromagnetic markers sensitive to the capacity limits of visual short-term memory. Frontiers in Human Neuroscience, 5(FEBRUARY), 1–20. 10.3389/fnhum.2011.00018

Naughtin, C. K., Mattingley, J. B., & Dux, P. E. (2016). Distributed and Overlapping Neural Substrates for Object Individuation and Identification in Visual Short-Term Memory. Cerebral Cortex, 26(2), 566–575. 10.1093/cercor/bhu212

Ngiam, W. X. Q. (2023). Mapping visual working memory models to a theoretical framework. Psychonomic Bulletin and Review, 0123456789. 10.3758/s13423-023-02356-5

Olson, I. R., & Berryhill, M. (2009). Some surprising findings on the involvement of the parietal lobe in human memory. Neurobiology of Learning and Memory, 91(2), 155–165. 10.1016/j.nlm.2008.09.006

Pylyshyn, Z. W. (1989). The role of location indexes in spatial perception. Cognition, 32, 65–97.

Robitaille, N., Grimault, S., & Jolicœur, P. (2009). Bilateral parietal and contralateral responses during maintenance of unilaterally encoded objects in visual short-term memory: Evidence from magnetoencephalography. Psychophysiology, 46(5), 1090–1099. 10.1111/j.1469-8986.2009.00837.x

Robitaille, N., Marois, R., Todd, J., Grimault, S., Cheyne, D., & Jolicœur, P. (2010). Distinguishing between lateralized and nonlateralized brain activity associated with visual short-term memory: fMRI, MEG, and EEG evidence from the same observers. NeuroImage, 53(4), 1334–1345. 10.1016/j.neuroimage.2010.07.027

Rouder, J. N., Morey, R. D., Morey, C. C., & Cowan, N. (2011). How to measure working memory capacity in the change detection paradigm. Psychonomic Bulletin and Review, 18(2), 324–330. 10.3758/s13423-011-0055-3

Swan, G., & Wyble, B. (2014). The binding pool: A model of shared neural resources for distinct items in visual working memory. Attention, Perception, and Psychophysics, 76(7), 2136–2157. 10.3758/s13414-014-0633-3

Taulu, S., & Simola, J. (2006). Spatiotemporal signal space separation method for rejecting nearby interference in MEG measurements. Physics in Medicine and Biology, 51(7), 1759–1768. 10.1088/0031-9155/51/7/008

Thyer, W., Adam, K. C. S., Diaz, G. K., Velázquez Sánchez, I. N., Vogel, E. K., & Awh, E. (2022). Storage in Visual Working Memory Recruits a Content-Independent Pointer System. Psychological Science, 33(10), 1680–1694. 10.1177/09567976221090923

Todd, J. J., & Marois, R. (2005). Posterior parietal cortex activity predicts individual differences in visual short-term memory capacity. Cognitive, Affective, & Behavioral Neuroscience, 5(2), 144–155. 10.3758/CABN.5.2.144

van Doorn, J., van den Bergh, D., Bohm, U., Dablander, F., Derks, K., Draws, T., Evans, N., Gronau, Q. F., Hinne, M., Kucharský, Š., Ly, A., Marsman, M., Matzke, D., Raj, A., Sarafoglou, A., Stefan, A., Voelkel, J., & Wagenmakers, E.-J. (2019). The JASP Guidelines for Conducting and Reporting a Bayesian Analysis. 10.31234/osf.io/yqxfr

Vogel, E. K., & Machizawa, M. G. (2004). Neural activity predicts individual differences in visual working memory capacity. Nature, 428(6984), 748–751. 10.1038/nature02447

Wang, B., Cao, X., Theeuwes, J., Olivers, C. N. L., & Wang, Z. (2017). Separate capacities for storing different features in visual working memory. Journal of Experimental Psychology: Learning Memory and Cognition, 43(2), 226–236. 10.1037/xlm0000295

Wilson, K. E., Adamo, M., Barense, M. D., & Ferber, S. (2012). To bind or not to bind: Addressing the question of object representation in visual short-term memory. Journal of Vision, 12(8), 1–16. 10.1167/12.8.14

Xu, Y. (2017). Reevaluating the Sensory Account of Visual Working Memory Storage. Trends in Cognitive Sciences, 21(10), 794–815. 10.1016/j.tics.2017.06.013

Xu, Y. (2023). Parietal-driven visual working memory representation in occipito-temporal cortex. Current Biology, 33(20), 4516-4523.e5. 10.1016/j.cub.2023.08.080

Xu, Y., & Chun, M. M. (2006). Dissociable neural mechanisms supporting visual short-term memory for objects. Nature, 440(7080), 91–95. 10.1038/nature04262

Yu, X., & Lau, E. (2023). The Binding Problem 2.0: Beyond Perceptual Features. Cognitive Science, 47(2). 10.1111/cogs.13244

Zhang, W., & Luck, S. J. (2008). Discrete fixed-resolution representations in visual working memory. Nature, 453(7192), 233–235. 10.1038/nature06860

